# Investigation of the effect of temperature on the structure of SARS-Cov-2 Spike Protein by Molecular Dynamics Simulations

**DOI:** 10.1101/2020.06.10.145086

**Authors:** Soumya Lipsa Rath, Kishant Kumar

**Affiliations:** Department of Biotechnology, National Institute of Technology Warangal (NIT W), Telangana, India, 506004; Department of Chemical Engineering, National Institute of Technology Warangal (NIT W), Telangana, India, 506004

**Keywords:** Structural protein, receptor binding motif, N-Terminal Domain, Closed conformation, Temperature-sensitive

## Abstract

Statistical and epidemiological data imply temperature sensitivity of the SARS-CoV-2 coronavirus. However, the molecular level understanding of the virus structure at different temperature is still not clear. Spike protein is the outermost structural protein of the SARS-CoV-2 virus which interacts with the Angiotensin Converting Enzyme 2 (ACE2), a human receptor, and enters the respiratory system. In this study, we performed an all atom molecular dynamics simulation to study the effect of temperature on the structure of the Spike protein. After 200ns of simulation at different temperatures, we came across some interesting phenomena exhibited by the protein. We found that the solvent exposed domain of Spike protein, namely S1, is more mobile than the transmembrane domain, S2. Structural studies implied the presence of several charged residues on the surface of N-terminal Domain of S1 which are optimally oriented at 10-30 °C. Bioinformatics analyses indicated that it is capable of binding to other human receptors and should not be disregarded. Additionally, we found that receptor binding motif (RBM), present on the receptor binding domain (RBD) of S1, begins to close around temperature of 40 °C and attains a completely closed conformation at 50 °C. The closed conformation disables its ability to bind to ACE2, due to the burying of its receptor binding residues. Our results clearly show that there are active and inactive states of the protein at different temperatures. This would not only prove beneficial for understanding the fundamental nature of the virus, but would be also useful in the development of vaccines and therapeutics.

**Graphical Abstract:** 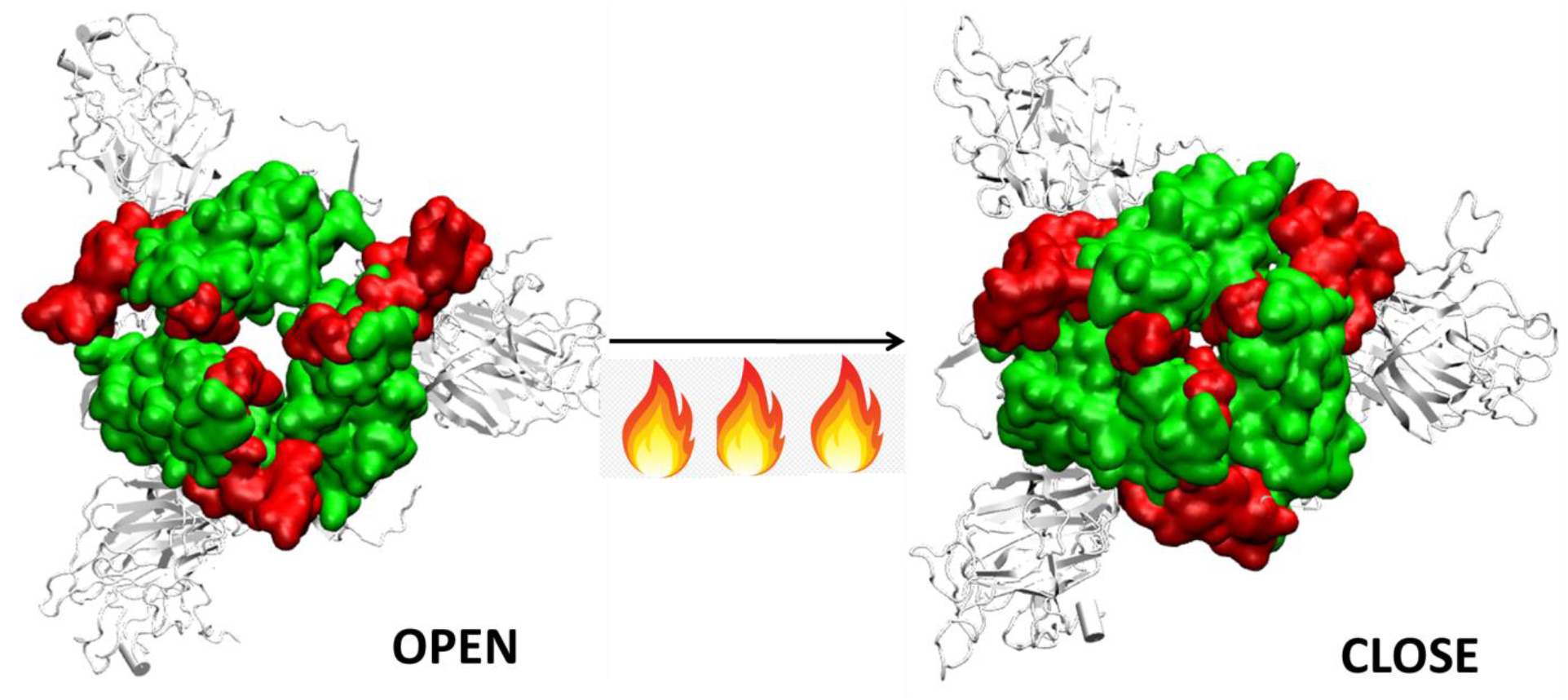

**Highlights:**

- Statistical and epidemiological evidence show that external climatic conditions influence the SARS-CoV infectivity, but we still lack a molecular level understanding of the same.
- Here, we study the influence of temperature on the structure of the Spike glycoprotein, the outermost structural protein, of the virus which binds to the human receptor ACE2.
- Results show that the Spike’s S1 domain is very sensitive to external atmospheric conditions compared to the S2 transmembrane domain.
- The N-terminal domain comprises of several solvent exposed charged residues that are capable of binding to human proteins. The region is specifically stable at temperatures ranging around 10-30° C.
- The Receptor Binding Motif adopts a closed conformation at 40°C and completely closes at higher temperatures making it unsuitable of binding to human receptors

## INTRODUCTION

Severe Acute Respiratory Syndrome Coronavirus 2 or SARS-COV-2, attacks the cells of the human respiratory system. Recent studies have found that the virus also interacts with the cells of the digestive system, renal system, liver, pancreas, eyes and brain [1]. It is known to cause severe sickness and is fatal in many cases [2]. It is believed that the virus originated in bats, which act as the natural reservoir; subsequently it got transmitted to human. It then gradually spread across almost all the nations through aerial transmission resulting in one of the worst known global pandemic of this century [3].

SARS-COV-2 is one of the seven forms of coronaviruses that affect the human population. The other known coronaviruses include HCoV-229E, HCoV-OC43, SARS-CoV, HCoV-NL63, HCoV-HKU1 and MERS-CoV [4,5]. Their infection varies from common cold to SARS, MERS or Covid19 [5]. These viruses have been observed to affect the human population predominantly during a particular season. For instance, the 2002 SARS infections began during the cold winters of November and after eight months, the number of reported cases became almost negligible [5]. Statistics show that countries with hot and humid weather conditions had lesser number of infectious cases of SARS [6]. However, MERS-COV, which was identified in Middle East regions, affected individuals during the summer [5]. Thus, the disease epidemiology suggests that the virus is found to be prominent in certain climatic conditions only.

The viability of SARS-COV-2 was measured on different surfaces by Chin et. al., who found that the virus droplets survived at 4 °C but quickly deactivated at elevated temperatures of 50 °C [7]. Smooth surfaces, plastics and iron show greater viability of the virus compared to that of paper, tissue, wood or cloth. Surgical masks had detectable viruses even on 7^th^ day [8, 9]. Soaps and disinfectants which disintegrate the virus membrane and structural proteins are a potent example of how the modulation of atmospheric conditions can affect the virus viability. Statistical reports by Cai et. al., and several others had shown that tropical countries like Malaysia, Indonesia or Thailand with high temperature and high relative humidity did not have major community outbreaks of SARS [6, 10–11]. Although viruses cannot be killed like bacteria by autoclaving, temperature sensitivity of virus have been reported several times in the past. Seasonal Rhinoviruses could not replicate at 37 °C whereas 33-35 °C is ideal for their survival in nasal cavity [12]. Influenza was found to be effective at a temperature around 37 °C, whereas higher temperatures of 41°C resulted in clumping of viruses on cell surfaces [13, 14, 15]. Similarly, the viability of SARS virus that persisted for 5 days at temperatures ranging between 22-25 °C and 40-50% humidity, was lost when the temperature was raised to 38 °C and 95% humidity [6].

When the virus is exposed to different temperature conditions, the initial interactions of the atmosphere occur with the structural proteins. There are four major structural proteins present on the virus, the Spike glycoprotein, the Envelope protein, the Membrane protein and the Nucleocapsid. Each of the proteins performs specific functions in receptor binding, viral assembly and genome release [16]. One of the first and largest structural proteins of the Coronavirus is the Spike glycoprotein [17]. The protein exists as a homotrimer where each monomer consists of 1273 amino acid residues (Figure 1) and is intertwined with each other. Each monomer has two domains, namely S1 and S2 [18]. The S1 and S2 domains are cleaved at a furin site by a host cell protease [18, 19]. The S1 domain lies predominantly above the lipid bilayer. The S2 domain, which is a class I transmembrane domain, travels across the bilayer and ends towards the inner side of the lipid membrane [18]. Figure 1 shows the two domains of the Spike glycoprotein.

**Figure 1:**
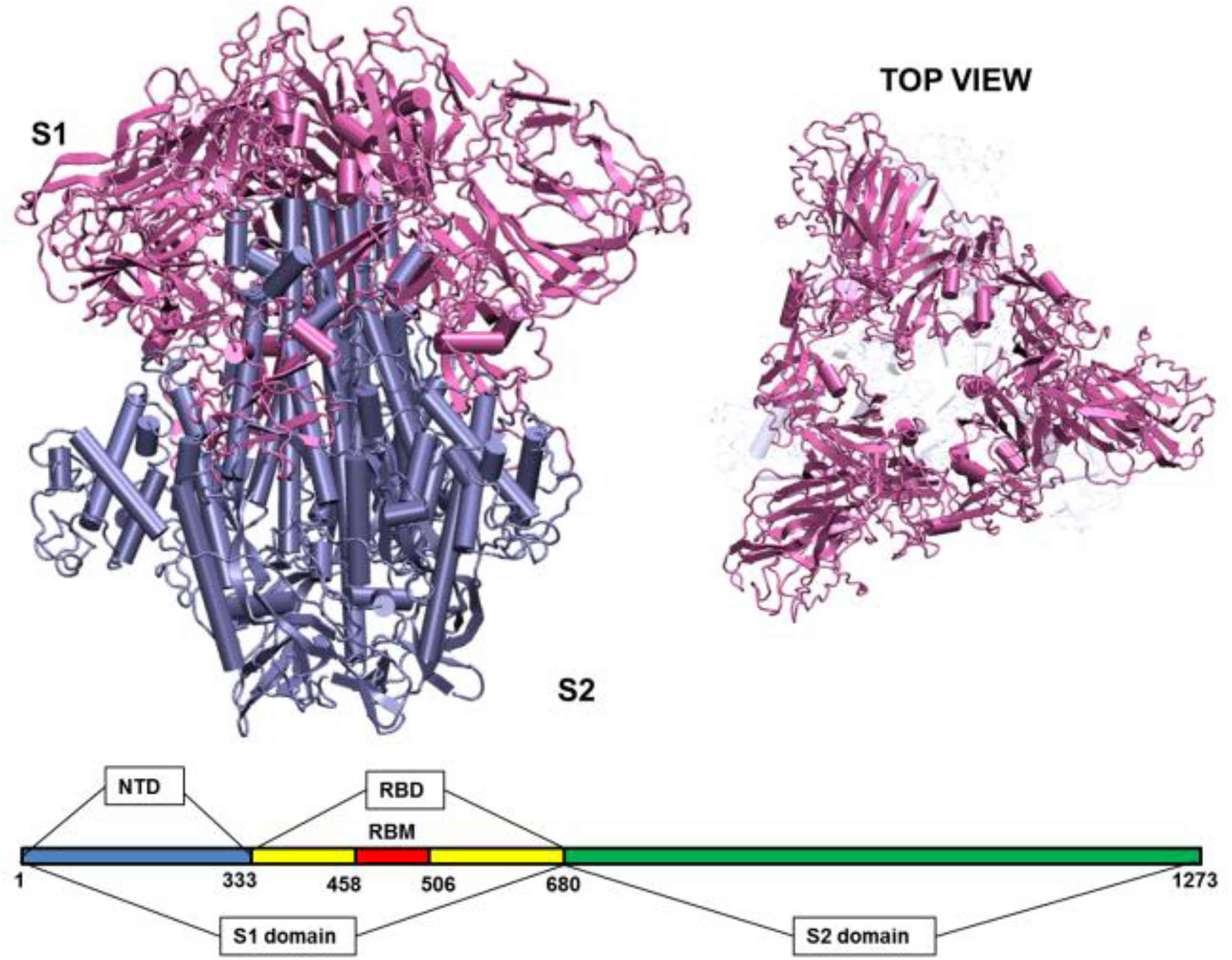
Structure of the Spike glycoprotein. The Spike glycoprotein, S1 and S2 domains, in the absence of glycan residues and lipid bilayer. The S1 domain is shown in pink and S2 in iceblue color. The top view of the protein shows a triangular arrangement of the S1 domain. Below the structure is a schematic showing the location of important regions on the S1 and S2 domains of the protein. The abbreviations in the S1 and S2 domains are: - NTD- N-Terminal Domain; RBD- Receptor Binding Domain; RBM- Receptor Binding Motif. The starting and ending residues are numbered.

The S1 domain comprises of mostly beta pleated sheets. It can be further classified into Receptor Binding domain (RBD) and N-terminal Domain (NTD). The RBD binds to Angiotensin Converting Enzyme 2 (ACE 2) on the host cells [20]. It lies on the top of the complex, where around 14 residues from the RBD domain bind to the ACE2 receptor on the host protein [21, 22]. The NTD is the outermost domain that is relatively more exposed and lies on the three sides giving a triangular shape to the protein when viewed from top (Figure 1). The NTD has a galectin fold and is known to bind to the sugar moieties [21]. The S2 domain on the other hand is a transmembrane region with strong interchain bonding between the residues. It is mostly α-helical and forms a triangle when viewed from bottom, though there is no overlapping of the top and bottom triangles.

Temperature is a very significant variable parameter for proteins because proteins respond differently in high and low temperature conditions. Many proteins have high thermal stability while others can unfold or even denature at high temperatures [23, 24]. During November, 2019, when the first outbreak of Covid19 was reported, the temperature in Wuhan, China was around 17 °C in the morning and 8°C at night. Tropical countries such as India, where a large number of cases still persist, had over 40 °C of temperature [25, 26]. Although statistical and experimental evidence show that temperature influences the activity and virulence of the virus, we still lack the understanding of the molecular level changes that are taking place in the virus due to the different weather conditions. Till date, there is no concrete evidence on whether atmospheric conditions actually influence the structure of the virus.

Here, by using all atom molecular dynamics (MD) simulations we explore the dynamics of the Spike glycoprotein of SARS-COV-2 at different temperatures. This is the first molecular study on the environmental influence on the protein structure. Results suggest that S1 domain is more flexible than S2. In the S1 domain, we observed the sensitivity of the receptor binding motif to different temperatures. We also found that the N-terminal domain of the protein has the potential of binding to different human receptors. The study will not only help us in understanding the nature of the virus but is also useful to design effective therapeutic strategies.

## RESULTS AND DISCUSSION

The crystal structure of the Spike glycoprotein (PDB: 6VXX) was found to have 871 missing residues. Thus, for our study we considered the complete model of the trimeric Spike protein generated by Zhang et. al. and had a Template modeling score of 0.6 [27]. The model was devoid of N-acetyl glucosamine (NAG) sugar moieties which are known to bind and stabilize the protein. The envelope lipid bilayer was not considered in the work to avoid large system size in atomistic simulations. After initial minimization and equilibration, we generated five different systems having temperatures ranging from 10°C to 50°C at an interval of 10 degrees. This was done to maintain the uniformity of the simulations, where temperature was the only variable that was different. In addition a temperature of 70°C was also imposed on the system to observe any possible deformation in the structure of spike protein, although this high temperature is not realistic to imitate the environmental condition (Table S1). Production run for 200ns was carried out in isothermal isobaric (NPT) ensemble.

### A. Spike glycoproteins are sensitive to temperature

After performing 200ns of classical Molecular dynamics simulations, the root mean square deviation (RMSD) of the trajectory, with respect to the starting structure, was calculated to check if the systems have attained stability. Figure S1 shows the complete RMSD of all the systems at different temperatures. It can be seen that the stability was attained within the first 50ns of the simulation time, thus, indicating that the systems are well equilibrated. The RMS values lie between 0.6 - 0.7nm for all the systems with an exception at 40°C where a marginally higher RMSD was seen after 100ns of simulation time. At temperatures 20°C and 30°C, a small rise in RMSD curves after 100ns of simulation time was observed. This implies that the Spike protein was more stable at temperatures 10°C and 50°C.

Since, the protein comprises of two distinct domains S1 and S2, we checked the RMSD of S1 and S2 domains individually, with respect to the starting structure, to understand the cause for higher RMSD values observed at 20°C, 30°C and 40°C (Figure 2). The RMS values of S1 domain at 20°C, 30°C and 40 °C were found to be around 0.7 nm, nearly 0.5nm more than simulations at 10 and 50 °C respectively. A similar trend was observed in the RMSD of S2 domain, but, the difference in values was only 0.15 nm. Although, in this study, we haven’t considered the bilayer lipid membrane of the SARS-COV-2 envelope inside which the Spike glycoprotein resides, the S2 domain shows remarkable stability in its RMSD values (Figure 2). The stability of the S2 domain can be conferred to the strong interchain interactions among the highly α-helical S2 domain.

**Figure 2:**
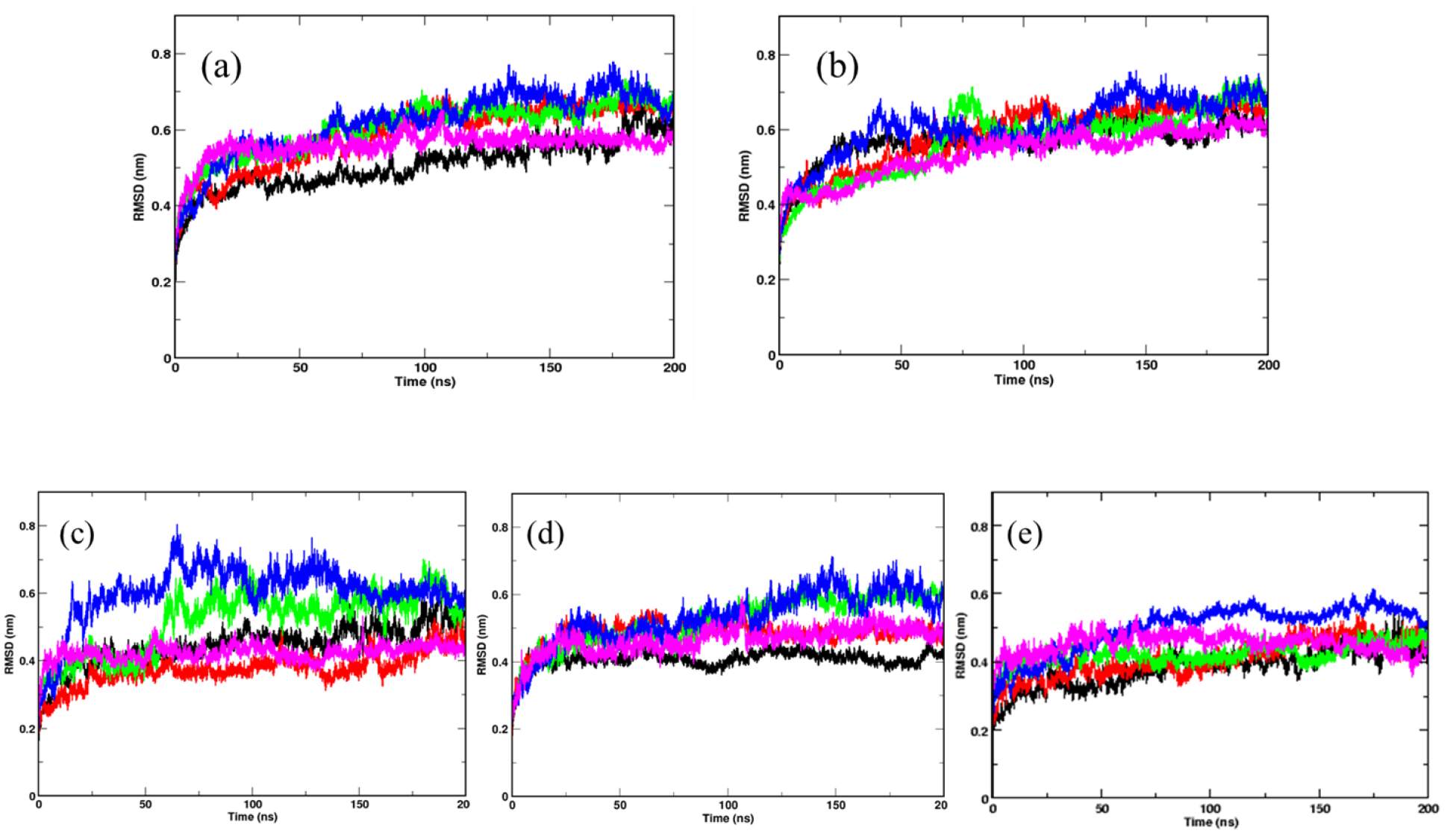
Temperature sensitivity of the S1 domain of Spike protein. RMSDs of Spike glycoprotein (a) S1 and (b) S2 domain showing stability of the S2 chains. The differential fluctuations of chains (c) A, (d) B and (e) C of S1 domain at 10 °C(black), 20 °C (red), 30 °C (green), 40 °C (blue) and 50 °C (magenta) implying effect of temperature of the chain stability.

Since the Spike protein is a homotrimer, the S1 domain of individual domains was also checked to account for the difference in fluctuations. Figure 2(c) - (e) shows the RMSD of S1 domain of chains A, B and C at different temperatures. In chain A, it can be clearly seen how the RMSD is quite high at temperatures of 30 °C and 40 °C respectively. At 50 °C however, the fluctuations are quite negligible and the system is very stable. Similarly, for chain B at 10 °C, 20 °C and 50 °C, the chains were stable. In the S1 domain of chain C, except for simulation at 40 °C, at all other temperatures, the RMSD values were found to be quite low along the length of the simulation time. The above data indicates that the protein chains, especially the S1 domains are quite flexible around the temperatures of 20-40 °C in comparison to low temperatures of 10 °C or high 50 °C of simulation temperature. Irrespective of the presence of the bilayer membrane, at different temperature conditions, the stalk of the Spike protein remains stable.

### B. Domain flexibility of S1 is more pronounced

In order to identify the region on the Spike protein that causes the deviations in RMSDs, we plotted the root mean square fluctuation (RMSF) of CA atoms of both S1 and S2 domains separately (Figure 3) at different temperatures. Each plot shows the RMSF of each individual chain at different temperatures. The RMSF of individual chains of S1 domain at different temperatures show that the residues ranging from 1-333 which constitute the N-Terminal Domain (NTD) of S1, show more fluctuations compared to the Receptor binding Domain (RBD) ranging from residues 334-680. In the NTD, three distinct peaks could be seen, viz:- residues 85-90, 100-200 and 240-260. The first peak in the NTD was observed around residues 85-90 (β4-β5), which is a loop directed inwards to the S2 domain (Figure S2). The peak was found to be highest in chain A at 40 °C (~0.8 nm), however at other temperatures all the chains have approximately 0.5 nm RMS fluctuation of its CA atoms. The residues 100-200 constitute the solvent exposed β sheet (β6-β12) of the NTD of S1 domain (Figure S2). The crystal structure (PDB: 6VXX) had shown as many as three glycosylated groups adjacent to this region of the protein (Figure S3) [28]. Thus, the lack of stabilizing sugar moieties in the simulated complex might play a role in increasing the flexibility of the region around 100-200. The residues 240-260 are solvent exposed loop around β14-β15. No glycan binding sites were observed in the crystal structure. The RBD domain consists of a receptor binding motif (RBM) ranging from residues 458-506 that show flexibility in all the systems. The lowest flexibility was observed at 10 °C. At 30 °C, the peaks were found for a wider range of residues. This indicates differential flexibility of the RBM at different temperatures. Since, the RBM is involved directly in binding to the ACE2 human receptor, its altered behavior at different temperatures would affect the protein-protein interaction.

**Figure 3:**
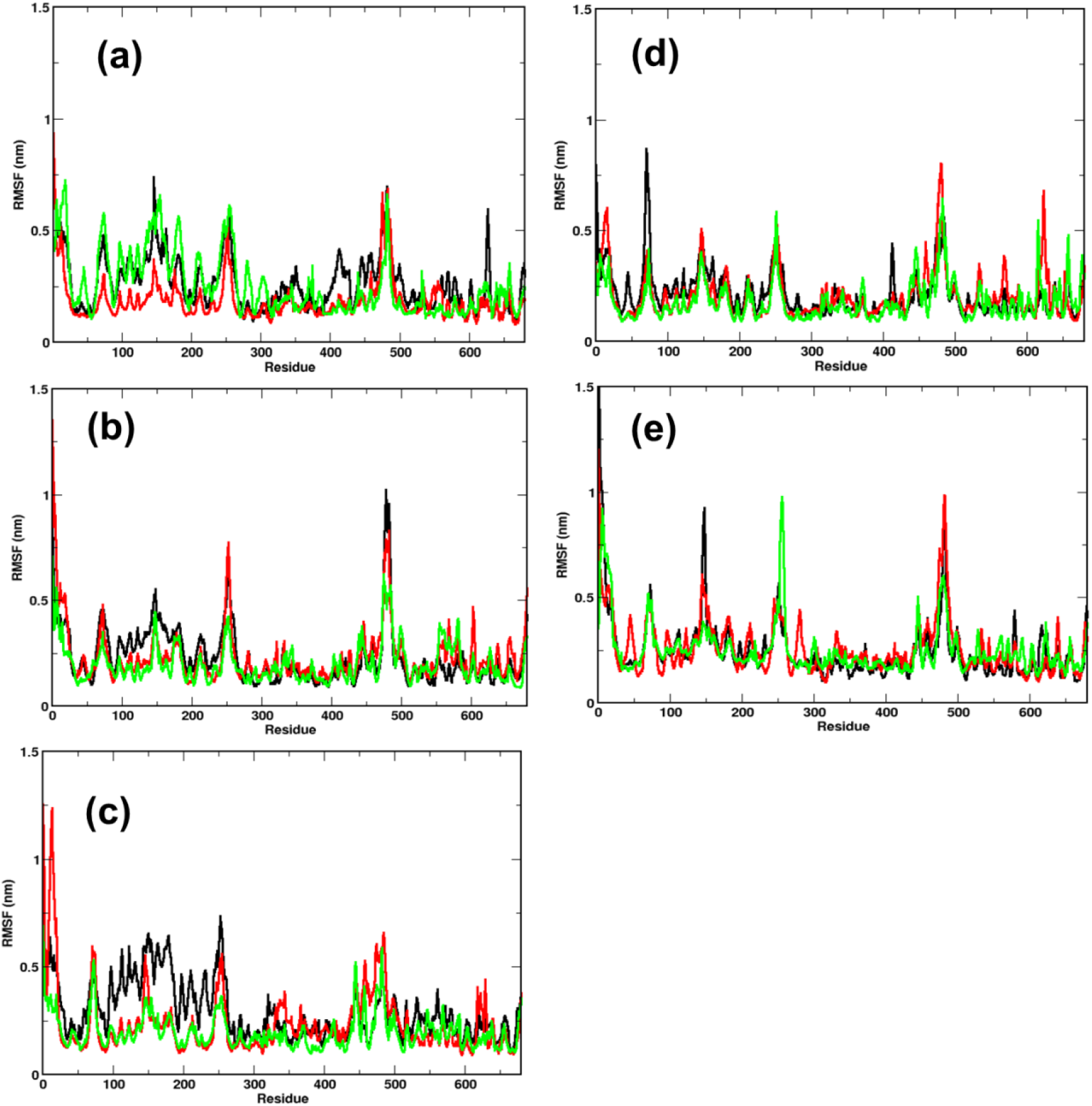
Fluctuation of CA of individual chains at different temperatures. RMSF of the CA atoms of the S1 domain for chains A (in black), B (in red) and C (in green) is shown. At temperatures a 10 °C, (b), 20 °C, (c) 30 °C, (d) 40 °C and (e) 50 °C, N-terminal Domain (residues 1-333) have higher mobility than the receptor binding domain (residues 334-680)

The RMSF of S2 domain on the other hand shows marked stability compared to domain S1 (Figure S4). This is in good agreement to our earlier observations of the RMSD of the S2 domain. Since it is a triple helical coil, the coiled-coil motif of the S2 domain which is further supported by three shorter helices supports domain stability [29]. However, the C-terminal residues 1125-1273 show greater flexibility compared to the rest of the domain. It should be noted that the C-terminal region of the Spike glycoprotein is exposed towards the inner side of the envelope bilayer and does not participate in the interchain interactions. It also has a more relaxed packing compared to the rest of the S2 [28, 30].

### C. NTD of the Spike protein could act as a receptor binding site

Although, the NTD is not known to directly bind to the receptor, in Mouse hepatitis coronavirus, it was found that the NTD binds to a CEACAM1a receptor [31]. Similarly, vaccines developed against the NTD of Spike protein in mice, had shown that NTD can also be a potential therapeutic target [32, 33]. Moreover, comparison between Bovine coronavirus and Bovine hemagglutinin-esterase enzyme indicated close evolutionary link between the virus and the host proteins, which could facilitate attachment in the host cells [34]. We performed a Multiple Sequence Alignment (MSA) of the SARS-CoV-2 coronavirus NTD with the human proteome to find similarity of NTD with human proteins

Figure 4 shows the phylogenetic tree based on fast minimum evolution. The tree was constructed from the results of protein Blast (Blastp) where the maximum allowed difference was 0.85 (details discussed in Materials and Methods). Results show similarity between the NTD and Briakunumab antibodies, anti-TSLP antibodies, Ephrin receptors, GABAA receptor, Leucine rich repeat containing Protein 4c, Kremen A receptor protein and N-galactosaminyltransferase. The remarkable similarity of the NTD with several human antibodies and receptor proteins indicate the possibility of the domain to be able to bind to different proteins in human. For instance, the Ephrin receptors are one of the largest proteins of receptor tyrosine kinases which are involved in angiogenesis, retinogenic signaling, axon guiding and in the migration of the epithelial cells of the intestines [1]. The possibility of the NTD acting as a receptor binding domain cannot be ruled out completely. There is also an uncanny correlation between the prevailing literature where scientists claim the virus affecting different parts of the human body and the similarity of NTD with human proteins [1, 35].

**Figure 4:**
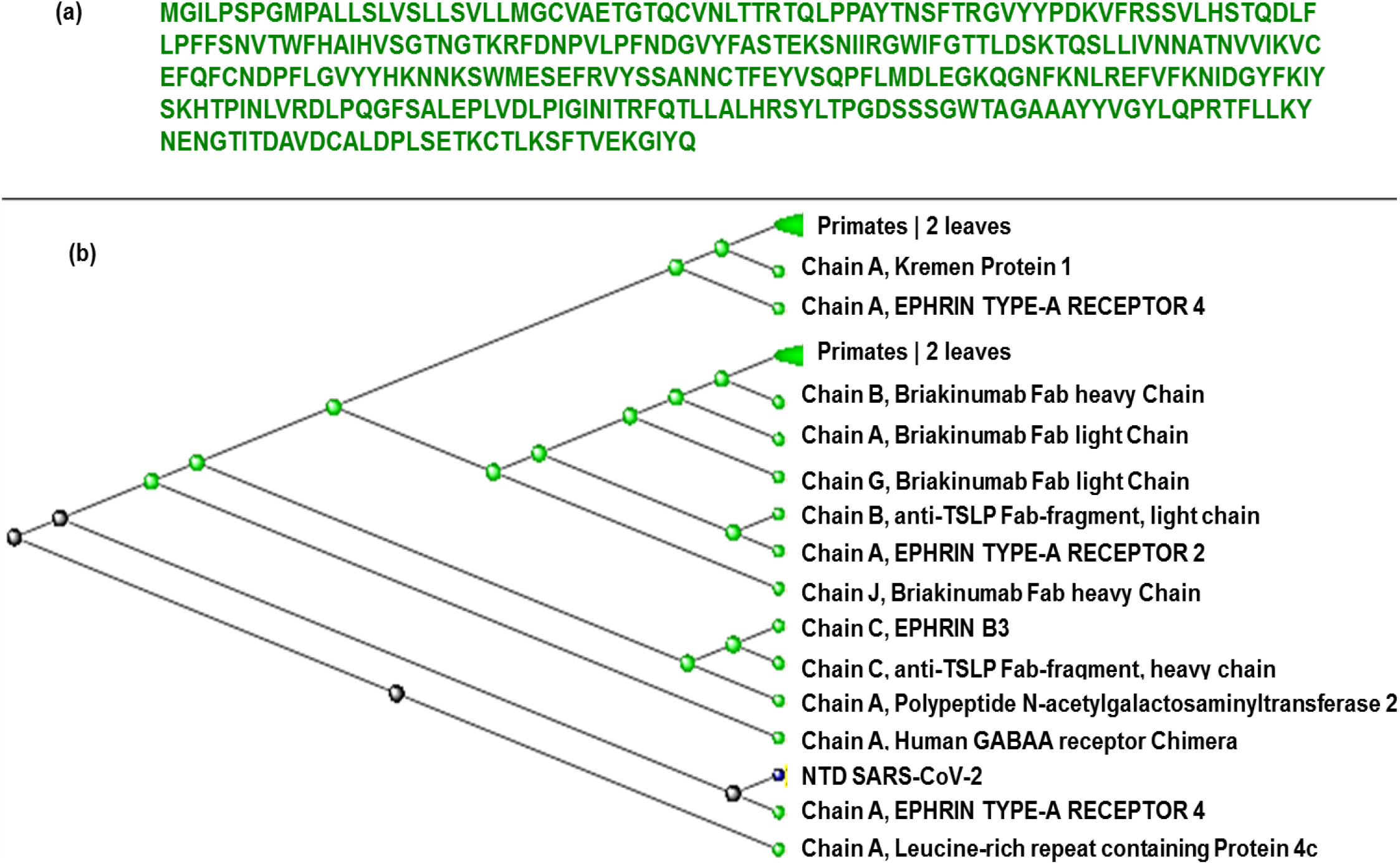
Similarity of NTD of SARS-CoV-2 with human proteins. (a) The sequence of the N-Terminal Domain of SARS-CoV-2 obtained from crystal structure which was used for Multiple sequence alignment and constructing phylogenetic tree (b) The Phylogenetic tree constructed from the results of multiple sequence alignments of N-Terminal domain of S1 and sequences of human proteome.

The NTD is relatively more exposed to solvents and more susceptible to external environmental conditions. However unlike RBD, the NTD doesn’t have a defined open or closed conformation. The coronavirus NTD is composed of three layered beta-sheet sandwich with 7, 3 and 6 antiparallel β strands in each layer making it a total of 16 beta stranded sheet with 5 prominent β hairpin loops (Figure S5). The crystal structures of Mouse Hepatitis Coronavirus (MHC) Spike protein and its receptor shows that the β1 and β6 of the NTD are the binding motif for CECAM1a protein [31]. However, unlike the MHC NTD, the arrangement of strands in SARS-CoV-2 is in opposite direction. The upper layer of the beta sandwich is composed of beta strands β4, β6, β7, β8, β9, β10, β14 (Figure S4). The three prominent regions which are exposed to the solvent and capable of interacting with potential receptors are regions N-terminal β strand, β8-β9, β9-β10 and β14-β15 loop.

Comparison of the NTD at different temperatures (Figure 5) show differential arrangement of the solvent exposed loops. The loops are formed by residues from N-terminal β strand, β8-β9, β9-β10, and β14-β15. The time averaged conformation of the loops after 200ns of simulation show that the loops are oriented close to each other at temperatures 10-30 °C, however at 40 °C and 50 °C, they move farther away from each other. Since, there was similarity between the Ehprin A proteins that binds to the Ephrin A receptors; we compared the residues involved in protein-protein interaction in the crystal structure of the human EphA4 ectodomain in complex with human Ephrin A5 for comparison. (PDB ID: 4BKA). There are three salt bridges and seven hydrogen bonds between the Ephrin protein and its receptor. Moreover, it can be clearly seen that the NTD loops host a large number of polar residues (Figure 5). These residues form a stable motif at temperatures 10-30 °C, primarily due to the stability between the loops. At 40 °C and 50 °C, hydrophobic patch from N-terminal β strand is exposed towards the solvent. The polar residues from β9-β10 and β14 −β15 move away from the N-terminal β strand and the β8-β9 loop, reducing the possibility of protein-protein interaction. Hence, a strong possibility exists for the NTD to act as a protein binding site at lower temperature ranges.

**Figure 5:**
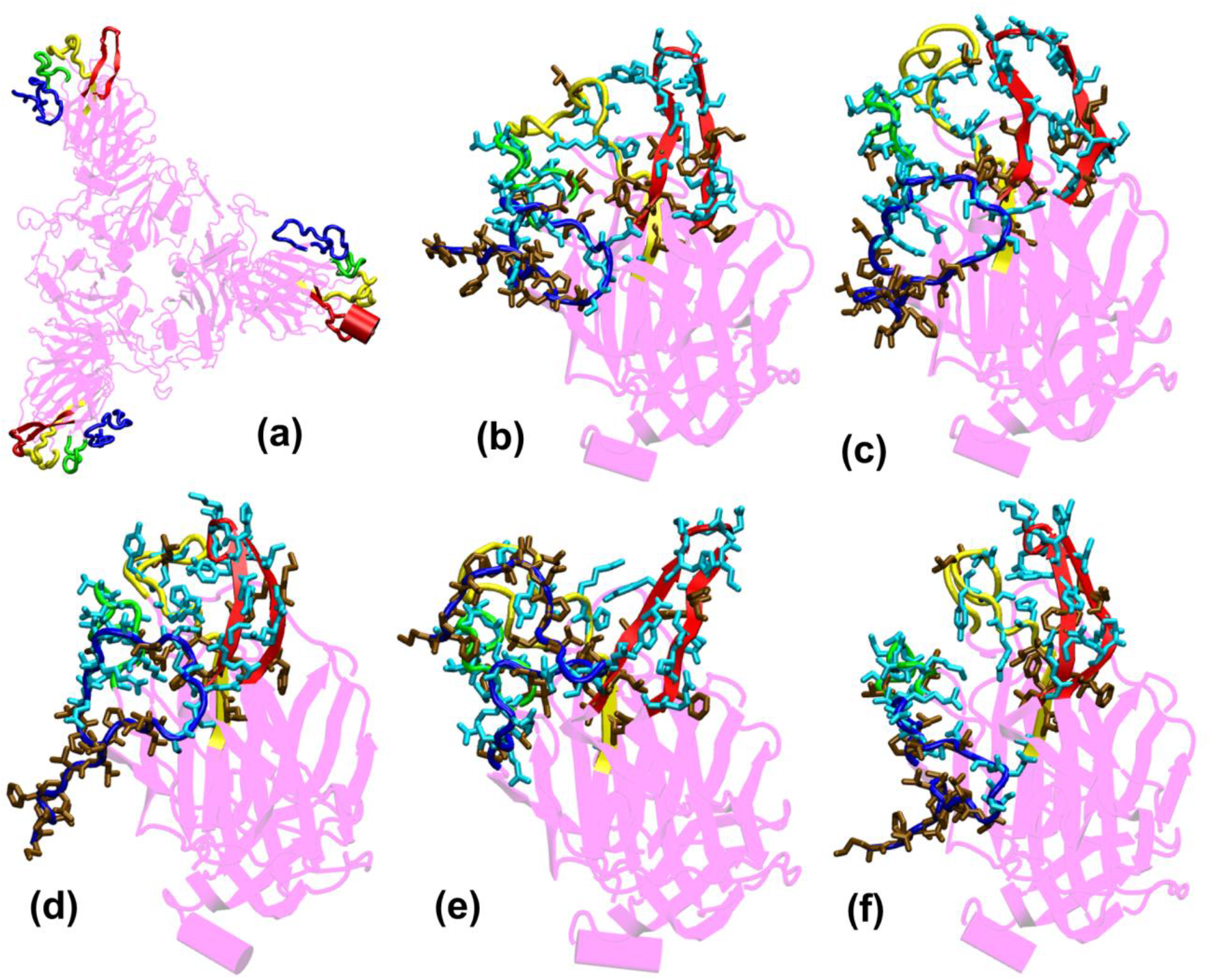
Structures of the N-Terminal Domain of Spike protein after 200ns of simulation showing the relative orientation of solvent exposed loops. (a) The solvent exposed loops of NTD; the N-terminal β strand, β8-β9, β9-β10 and β14-β15 are shown in red, blue, green and yellow colors respectively. Time-averaged conformation of N- Terminal Domain of SARS-CoV-2 Spike protein at, (b) 10 °C, (c) 20 °C, (d) 30 °C, (e) 40 °C and (f) 50 °C showing the relative orientation of the polar and hydrophobic residues. The residues are shown in licorice. Polar resides are colored in light blue and hydrophobic in brown colors respectively.

From the bioinformatics and structural analyses, we observed that the NTD not only acts as a glycan binding site but can also as a site for binding of several human proteins. The motif formed out of several polar residues on the solvent exposed loops at 10-30 °C could form salt-bridges and hydrogen bonds with partner proteins. At higher temperatures, the propensity of forming such interactions would be lost owing to the differential orientation of the loops. Nonetheless, the NTD could act as a possible target for development of vaccines and inhibitors.

### D. The RBD behaves differently at higher temperatures

The receptor binding domain (RBD) of the Spike glycoprotein is a potential target for vaccine and drug development [36, 37]. It is highly conserved among the human coronaviruses and binds to ACE2 receptor present on the lung tissues [38]. Residues 458-506 of the RBD domain comprises of the receptor binding motif (RBM). The RBM has 8 residues which are identical and 5 residues with similar biochemical properties between SARS, MERS and SARS-COV-2. This conserved region primarily interacts with the ACE2 receptor and hence, often scientists target the RBD domain of for developing therapeutic agents [36, 37, and 39]. Earlier in Figure 2, we saw that the RBD domain spanning from residues 333-680 shows higher stability when compared to the NTD of the S1 domain at different range of temperatures.

We compared the time averaged conformation of the RBD generated from the last 10ns of the simulation time at different temperatures (Figure 6). The core β pleated sheet was very stable demonstrating no lack of secondary structures at higher temperatures. However, the RBM motif (highlighted in magenta in Figure 6) shows a very dynamic conformation across different temperature ranges The dynamics was more pronouned at 10 °C, 20 °C and 30°C whereas at 40°C and 50°C of temperature, the RBM had a more confined conformation. The RBD flexibility was more apparent at 20°C and 30°C where the three chains moved further away from each other. However, a tighter and well packed structure was found for the protein at 50 °C. The figures suggest that although residue wise movements in RBD were not visible in RMSF (Figure 2), the RBD domains and motifs show intrinsic flexibility along particular temperature ranges. Previous studies have indicated that the RBD domain can adopt either an open or a closed conformation in the virus [18]. We compared the conformation of the Spikeprotein-ACE2 crystal structure and found that in the open conformation, the RBD exposes its RBM residues Phe456, Ala475, Phe486, Asn487, Tyr489, Gln493, Gly496, Gln498, Thr500, Asn501, Gly502 and Tyr505 to failitate the binding of the ACE2 receptors. It is fascinating to see that at 40 °C and more interestingly at 50°C, the RBM motif is in a closed loop conformation and very compact which hinders its association with the partner proteins.

**Figure 6:**
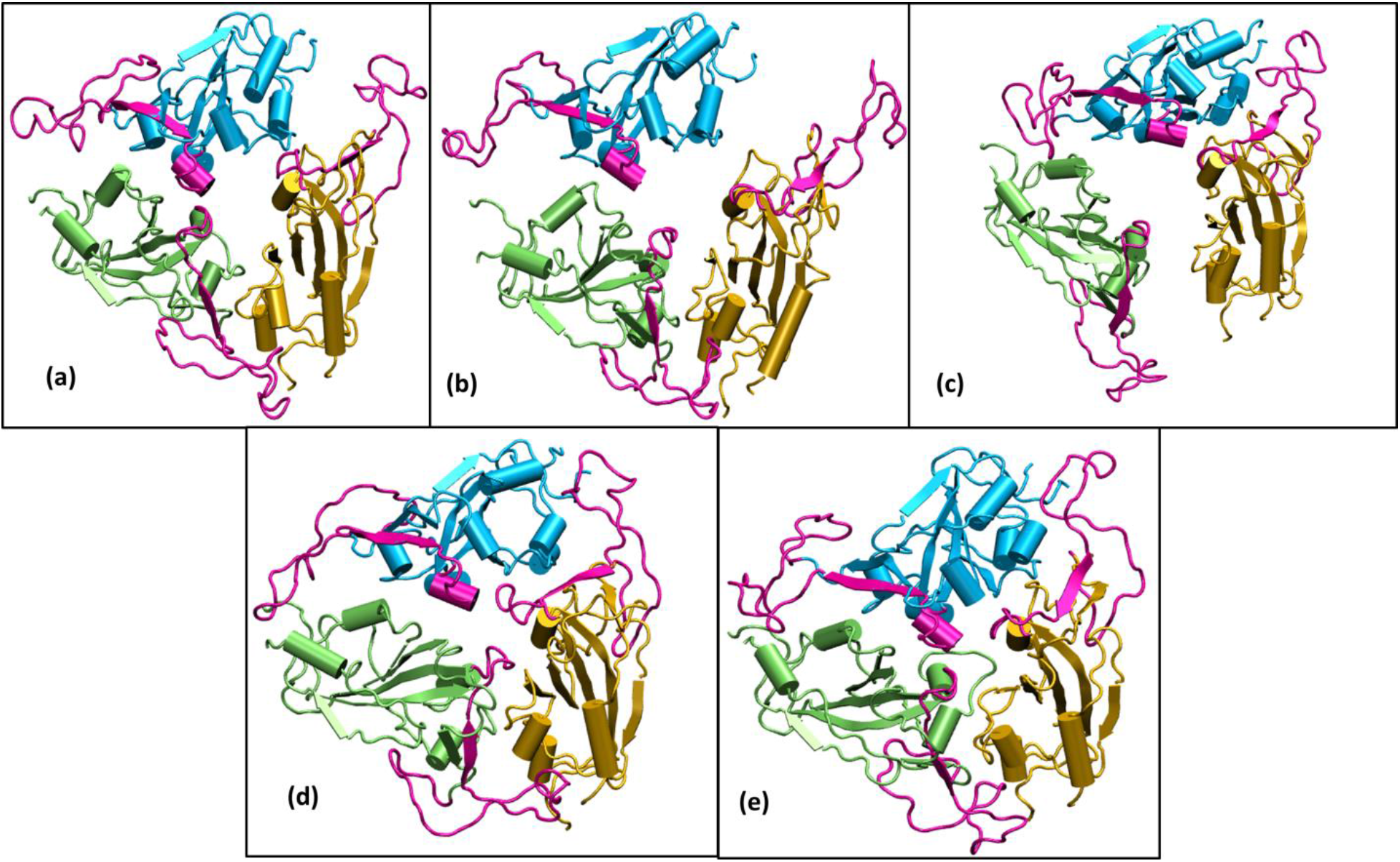
Structures of the receptor binding domain of Spike protein after 200ns of simulation at different temperatures exhibit diverse structural dynamics. Time-averaged conformations of RBD of SARS-CoV-2 Spike protein at, (a) 10 °C, (b) 20 °C, (c) 30 °C, (d) 40 °C and (e) 50 °C. The three chains are colored in lime, cyan and orange. The Receptor binding motif (shown in magenta) is oriented in a confined conformation at higher temperatures.

To validate the findings, we ran another simulation of the Spike protein at a higher temperature of 70 °C. After 100ns of simulation, we found that significant similarity between the closed conformation observed at 50 °C and the conformation at 70 °C. The RBM residues, specifically Phe456, Ala475, Phe486, Asn487, Tyr489, Gln493, Gly496, Gln498, Thr500, Asn501, Gly502 and Tyr505 were found to be clearly buried between the interchain subunits at 70 °C (Figure 7, S6). However, when compared to the orientation at 30 °C the residues are directed towards the solvent Thus the reason for very stable RMSD observed in Figure 1 is largely due to the confined architecture of the receptor binding domain at 50°C and higher temperatures. The unavailability of RBM residues to bind to ACE2 receptor would nonetheless destabilize virus-protein interactions at higher temperatures.

**Figure 7:**
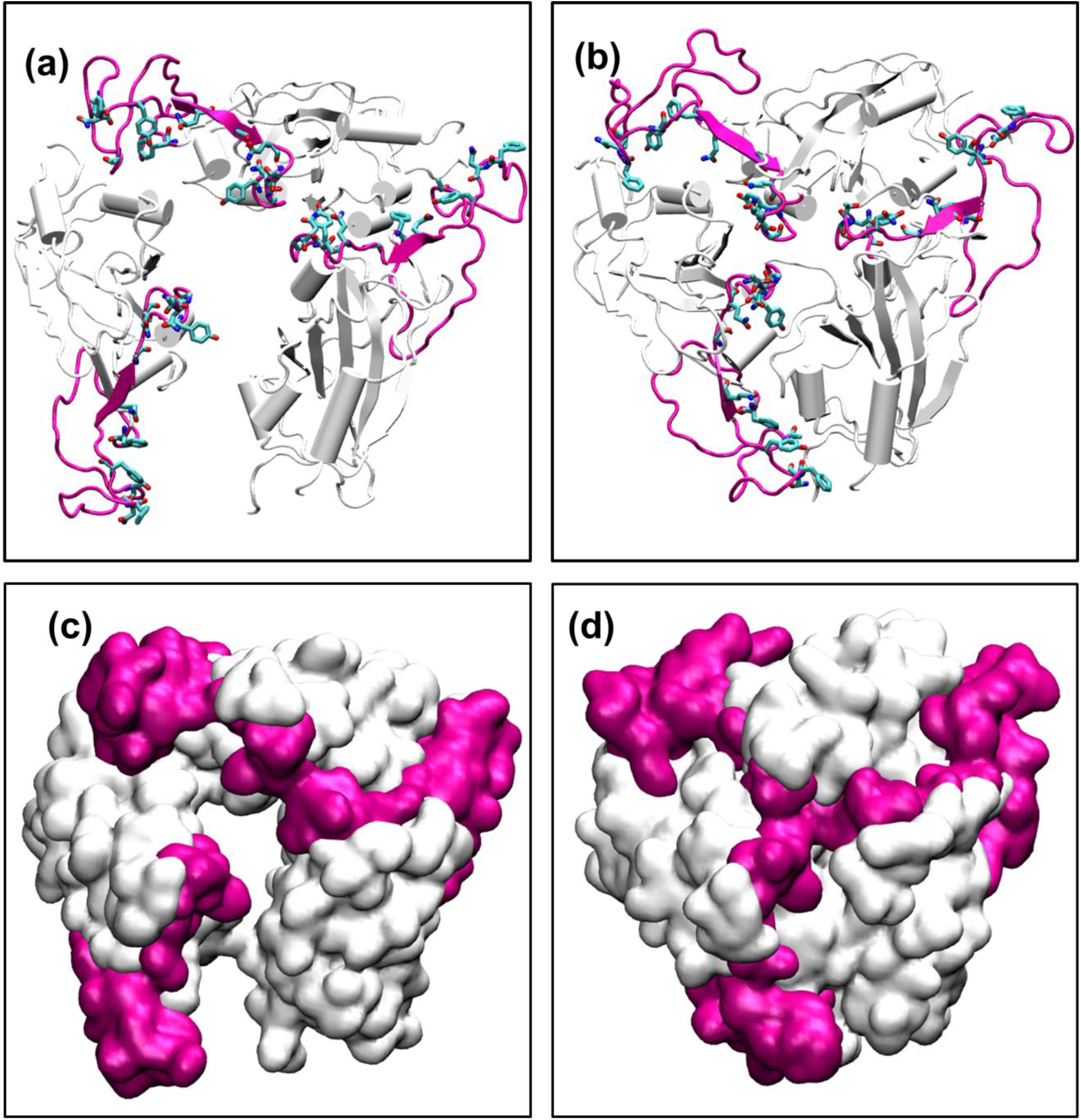
The conformations of Receptor Binding motif. The time averaged structures of the Spike protein showing the open and closed conformations of the receptor binding domain at (a) 30°C and (b) 70 °C. The residues on the Receptor binding motif (shown as sticks and colored by CPK) are buried at the interchain interface at high temperatures but readily exposes its residues at 30 °C. (c) and (d) show the open and closed conformations of the receptor binding domain at (a) 30°C and (b) 70 °C in surf mode. The receptor binding motif is colored in magenta and the receptor binding domain is shown in white.

Thus, although the RBM stays largely in open conformation state, surprisingly, from around the temperature of 40°C a closed conformation of the motif was observed. In this conformation, the RBM from the three chains come very close to each other sealing the visibility of the trimeric pore. At temperatures >50 °C, the Spike RBM is completely closed (Figure 7). The closing of the RBM buries the receptor binding residues inside trimer abolishing the possibility of contacts with the ACE2 receptor and making the Spike protein inactive. Our results clearly show that the activity of the Spike protein is dependent on the external temperatures where a higher temperature renders it completely inactive.

## CONCLUSION

The SARS-CoV-2 has severely affected the human population with large number of infected individuals around world. The propensity of virus to survive in cold and dry climatic conditions have been speculated by researchers and supported by the statistical evidence from earlier SARS epidemic of 2002. However, it is still unclear how the virus undergoes changes at the structural level in different environments. The Spike protein of the virus helps in the attachment and entry of the coronaviruses inside the host cells. It exists as a homotrimer and is partly exposed to the outer environment and partly immersed inside the lipid bilayer of the viral envelope. Here, we studied the differential response of the Spike protein at different temperature conditions.

Our results show that the S2 transmembrane domain remains stable even without the bilayer membrane, whereas the solvent exposed S1 domain is quite flexible. Moreover, the S1 comprises of two subdomains, namely N-terminal domain (NTD) and the receptor binding domain (RBD). The simulations results show that the RBD is relatively less mobile. Its flexibility is limited only to the receptor binding motif or RBM which interacts with the Angiotensin Converting enzyme 2 (ACE2), its human receptor. However, the NTD was found to be quite mobile.

Although, the NTD doesn’t directly interact with the ACE2 receptor in humans, it has been found to bind to receptors in other mammals [31]. The flexible NTD hosts a large number of charged residues on the top layer of its tri-layered beta sandwich architecture. However at 40-50°C of temperature, the polar residues were found to be less solvent exposed. The similarity of the NTD sequence with the several human receptors such as Ephrins, Briakunumab, anti-TSLP, etc indicated a possibility of the subdomain to be involved in binding to alternate human proteins.

The RBM present on the RBD is very crucial in initial protein-protein interaction between the host and virus. We found that this domain is largely in an open conformation which enables receptor binding at lower temperatures. Surprisingly, from ~40°C, a closed conformation of the motif was observed. In this conformation, the RBM from the three chains come very close to each other sealing the visibility of the trimeric pore. At temperatures >50°C, the Spike RBM is completely closed. The closing of the RBM buries the receptor binding residues inside trimer abolishing the possibility of contacts with the ACE2 receptor and making the Spike protein inactive.

Our results have shown for the first time that the Spike protein has the possibility to stay in an active and inactive state based on the external temperature. Since no visible loss of secondary structure was observed at higher temperatures, it would be interesting to know if the conformational change is reversible in nature. Nevertheless, this work would prove very beneficial in the development of vaccines as well as development of therapeutic strategies that target not only the receptor binding domain but also the N-Terminal Domain of the Spike protein.

## MATERIALS AND METHODS

The complete model of the Spike glycoprotein of SARS-COV-2 was obtained from Zhang lab (GenBank: QHD43416.1). This model was considered because it had modeled the missing 871 residues that were missing in the crystal structure 6VXX. It had a Template Modeling (TM) score of 0.6 [27]. The initial Root Mean Square Deviation (RMSD) between the model and the closed crystal structure of Spike glycoprotein (PDB ID 6VXX) was found to be 1.54 Å. The model was devoid of N-acetyl glycosamine (NAG) glycan residues and consisted of the glycoprotein trimer where each monomer had aminoacids ranging from 1 to 1273. The structure was initially solvated with a water box having a cubic box of size 17.9 x 17.9 x 17.9 nm and 569293 atoms with water and ions. The minimum distance between the protein and the edge of the water box was fixed at 13 Å. Particle-Mesh Ewald (PME) method was used for electrostatic interactions using a grids pacing of 0.16nm and a 1.0 nm cutoff. After energy minimization and equilibration, by maintaining harmonic restraints on the protein heavy atoms, the system was gradually heated to 300K in a canonical ensemble. The harmonic restraints were gradually reduced to zero and solvent density was adjusted under isobaric and isothermal conditions at 1 atm and 300 K. This was followed by 500ps NVT and 500ps NPT equilibration with harmonic restraints of 1000 kJ mol^−1^ nm^−2^ on the heavy atoms. Production run for all the systems was carried out for 200ns till it reached a stable RMSD. All simulations were carried out in Gromacs 2020 with AMBERff99SB-ILDN forcefield for proteins [40, 41]. The long-range electrostatic interactions were treated by using Particle-Mesh Ewald sum and SHAKE was used to constrain all bonds involving hydrogen atoms. After equilibration, systems were heated or cooled at different temperatures (Table S1) and simulated for 200ns. All analyses were carried out using Gromacs analysis tools [40]. Protein Blast was used to search similar sequences in the human proteome. The Blast Tree View widget helped us generate the phylogenetic tree which is a simple distance based clustering of the sequences based on pairwise alignment results of Blast relative to the query sequence [42]. VMD was used for visualization of results and generation of figures [43].

## Supporting information

Supoorting Information

## SUPPORTING INFORMATION

Supporting figures, Figs S1-S6 and Table S1 are provided online.

## ACKNOWLEDGEMENTS

This research used resources of the National Energy Research Scientific Computing Center of the Ernest Orlando Lawrence Berkeley National Laboratory, a DOE Office of Science User Facility supported by the Office of Science of the U.S. Department of Energy under Contract No. DE-AC02-05CH11231 and used the Extreme Science and Engineering Discovery Environment (XSEDE). The authors are thankful to the Covid19 HPC Consortium for providing resources and helping researchers work for a noble cause. The authors are also thankful to Dr. Suchetana Gupta, Dr. Debakanta Tripathy and Dr. Chockalingam S for critically proof reading the manuscript. We are also grateful to National Institute of Technology Warangal for providing facilities.

## Abbreviations

RBD: receptor binding domain
NTD: N-Terminal Domain
RBM: receptor binding motif
ACE2: Angiotensin Converting enzyme 2

## REFERENCES

[1] D.E. Gordon, G.M. Jang, M. Bouhaddou, J. Xu, K. Obernier, K.M. White, M.J. O’Meara, V.V. Rezelj, J.Z. Guo, D.L. Swaney, T.A. Tummino, R. Huettenhain, R.M. Kaake, A.L. Richards, B. Tutuncuoglu, H. Foussard, J. Batra, K. Haas, M. Modak, M. Kim, P. Haas, B.J. Polacco, H. Braberg, J.M. Fabius, M. Eckhardt, M. Soucheray, M.J. Bennett, M. Cakir, M.J. McGregor, Q. Li, B. Meyer, F. Roesch, T. Vallet, A. Mac Kain, L. Miorin, E. Moreno, Z.Z.C. Naing, Y. Zhou, S. Peng, Y. Shi, Z. Zhang, W. Shen, I.T. Kirby, J.E. Melnyk, J.S. Chorba, K. Lou, S.A. Dai, I. Barrio-Hernandez, D. Memon, C. Hernandez-Armenta, J. Lyu, C.J.P. Mathy, T. Perica, K.B. Pilla, S.J. Ganesan, D.J. Saltzberg, R. Rakesh, X. Liu, S.B. Rosenthal, L. Calviello, S. Venkataramanan, J. Liboy-Lugo, Y. Lin, X.P. Huang, Y.F. Liu, S.A. Wankowicz, M. Bohn, M. Safari, F.S. Ugur, C. Koh, N.S. Savar, Q.D. Tran, D. Shengjuler, S.J. Fletcher, M.C. O’Neal, Y. Cai, J.C.J. Chang, D.J. Broadhurst, S. Klippsten, P.P. Sharp, N.A. Wenzell, D. Kuzuoglu, H.Y. Wang, R. Trenker, J.M. Young, D.A. Cavero, J. Hiatt, T.L. Roth, U. Rathore, A. Subramanian, J. Noack, M. Hubert, R.M. Stroud, A.D. Frankel, O.S. Rosenberg, K.A. Verba, D.A. Agard, M. Ott, M. Emerman, N. Jura, M. von Zastrow, E. Verdin, A. Ashworth, O. Schwartz, C. d’Enfert, S. Mukherjee, M. Jacobson, H.S. Malik, D.G. Fujimori, T. Ideker, C.S. Craik, S.N. Floor, J.S. Fraser, J.D. Gross, A. Sali, B.L. Roth, D. Ruggero, J. Taunton, T. Kortemme, P. Beltrao, M. Vignuzzi, A. García-Sastre, K.M. Shokat, B.K. Shoichet, N.J. Krogan, A SARS-CoV-2 protein interaction map reveals targets for drug repurposing, Nature. (2020). https://doi.org/10.1038/s41586-020-2286-9.

[2] World Health Organization, 2020

[3] J.S. MacKenzie, D.W. Smith, COVID-19: A novel zoonotic disease caused by a coronavirus from China: What we know and what we don’t, Microbiol. Aust. 41 (2020) 45–50. https://doi.org/10.1071/MA20013.

[4] H. Gao, H. Yao, S. Yang, L. Li, From SARS to MERS: evidence and speculation, Front. Med. 10 (2016) 377–382. https://doi.org/10.1007/s11684-016-0466-7.

[5] S. Su, G. Wong, W. Shi, J. Liu, A.C.K. Lai, J. Zhou, W. Liu, Y. Bi, G.F. Gao, Epidemiology, Genetic Recombination, and Pathogenesis of Coronaviruses, Trends Microbiol. 24 (2016) 490–502. https://doi.org/10.1016/j.tim.2016.03.003.

[6] K.H. Chan, J.S.M. Peiris, S.Y. Lam, L.L.M. Poon, K.Y. Yuen, W.H. Seto, The effects of temperature and relative humidity on the viability of the SARS coronavirus, Adv. Virol. 2011 (2011). https://doi.org/10.1155/2011/734690.

[7] A.W.H. Chin, J.T.S. Chu, M.R.A. Perera, K.P.Y. Hui, H.-L. Yen, M.C.W. Chan, M. Peiris, L.L.M. Poon, Stability of SARS-CoV-2 in different environmental conditions, The Lancet Microbe. 1 (2020) e10. https://doi.org/10.1016/s2666-5247(20)30003-3.

[8] L.M. Casanova, S. Jeon, W.A. Rutala, D.J. Weber, M.D. Sobsey, Effects of air temperature and relative humidity on coronavirus survival on surfaces, Appl. Environ. Microbiol. 76 (2010) 2712–2717. https://doi.org/10.1128/AEM.02291-09.

[9] N. Van Doremalen, T. Bushmaker, D.H. Morris, M.G. Holbrook, A. Gamble, B.N. Williamson, A. Tamin, J.L. Harcourt, N.J. Thornburg, S.I. Gerber, J.O. Lloyd-Smith, E. De Wit, V.J. Munster, Aerosol and surface stability of SARS-CoV-2 as compared with SARS-CoV-1, N. Engl. J. Med. 382 (2020) 1564–1567. https://doi.org/10.1056/NEJMc2004973.

[10] Q.C. Cai, J. Lu, Q.F. Xu, Q. Guo, D.Z. Xu, Q.W. Sun, H. Yang, G.M. Zhao, Q.W. Jiang, Influence of meteorological factors and air pollution on the outbreak of severe acute respiratory syndrome, Public Health. 121 (2007) 258–265. https://doi.org/10.1016/j.puhe.2006.09.023.

[11] J. Tan, L. Mu, J. Huang, S. Yu, B. Chen, J. Yin, An initial investigation of the association between the SARS outbreak and weather: With the view of the environmental temperature and its variation, J. Epidemiol. Community Health. 59 (2005) 186–192. https://doi.org/10.1136/jech.2004.020180.

[12] E.F. Foxman, J.A. Storer, M.E. Fitzgerald, B.R. Wasik, L. Hou, H. Zhao, P.E. Turner, A.M. Pyle, A. Iwasaki, Temperature-dependent innate defense against the common cold virus limits viral replication at warm temperature in mouse airway cells, Proc. Natl. Acad. Sci. U. S. A. 112 (2015) 827–832. https://doi.org/10.1073/pnas.1411030112.

[13] A.C. Lowen, J. Steel, Roles of Humidity and Temperature in Shaping Influenza Seasonality, J. Virol. 88 (2014) 7692–7695. https://doi.org/10.1128/jvi.03544-13.

[14] Y.I. Ishida, A. Hiraki, E. Hirayama, Y. Koga, J. Kim, Temperature-sensitive viral infection: Inhibition of hemagglutinating virus of Japan (Sendai virus) infection at 41°, Intervirology. 45 (2002) 125–135. https://doi.org/10.1159/000065865.

[15] I. Pelletier, D. Rousset, V. Enouf, F. Colbère-Garapin, S. van der Werf, N. Naffakh nadianaffakh, Highly heterogeneous temperature sensitivity of 2009 pandemic influenza A(H1N1) viral isolates, northern France, 2011. www.eurosurveillance.org:pii=19999. Available online:http://www.eurosurveillance.org/ViewArticle.aspx?ArticleId=19999.

[16] I. Astuti, Ysrafil, Severe Acute Respiratory Syndrome Coronavirus 2 (SARS-CoV-2): An overview of viral structure and host response, Diabetes Metab. Syndr. Clin. Res. Rev. 14 (2020) 407–412. https://doi.org/10.1016/j.dsx.2020.04.020.

[17] F. Li, Structure, Function, and Evolution of Coronavirus Spike Proteins, Annu. Rev. Virol. 3 (2016) 237–261. https://doi.org/10.1146/annurev-virology-110615-042301.

[18] A.C. Walls, Y.J. Park, M.A. Tortorici, A. Wall, A.T. McGuire, D. Veesler, Structure, Function, and Antigenicity of the SARS-CoV-2 Spike Glycoprotein, Cell. 181 (2020) 281–292.e6. https://doi.org/10.1016/j.cell.2020.02.058.

[19] S. Belouzard, V.C. Chu, G.R. Whittaker, Activation of the SARS coronavirus spike protein via sequential proteolytic cleavage at two distinct sites, n.d. www.pnas.org/cgi/content/full/.

[20] Q. Wang, Y. Zhang, L. Wu, S. Niu, C. Song, Z. Zhang, G. Lu, C. Qiao, Y. Hu, K.Y. Yuen, Q. Wang, H. Zhou, J. Yan, J. Qi, Structural and Functional Basis of SARS-CoV-2 Entry by Using Human ACE2, Cell. 181 (2020) 894–904.e9. https://doi.org/10.1016/j.cell.2020.03.045.

[21] Y. Yuan, D. Cao, Y. Zhang, J. Ma, J. Qi, Q. Wang, G. Lu, Y. Wu, J. Yan, Y. Shi, X. Zhang, G.F. Gao, Cryo-EM structures of MERS-CoV and SARS-CoV spike glycoproteins reveal the dynamic receptor binding domains, Nat. Commun. 8 (2017). https://doi.org/10.1038/ncomms15092.

[22] J. Lan, J. Ge, J. Yu, S. Shan, H. Zhou, S. Fan, Q. Zhang, X. Shi, Q. Wang, L. Zhang, X. Wang, Structure of the SARS-CoV-2 spike receptor-binding domain bound to the ACE2 receptor, Nature. 581 (2020) 215–220. https://doi.org/10.1038/s41586-020-2180-5.

[23] L. Julió Plana, A.D. Nadra, D.A. Estrin, F.J. Luque, L. Capece, Thermal Stability of Globins: Implications of Flexibility and Heme Coordination Studied by Molecular Dynamics Simulations, J. Chem. Inf. Model. 59 (2019) 441–452. https://doi.org/10.1021/acs.jcim.8b00840.

[24] Y.W. Dong, M.L. Liao, X.L. Meng, G.N. Somero, Structural flexibility and protein adaptation to temperature: Molecular dynamics analysis of malate dehydrogenases of marine molluscs, Proc. Natl. Acad. Sci. U. S. A. 115 (2018) 1274–1279. https://doi.org/10.1073/pnas.1718910115.

[25] Y. Wu, W. Jing, J. Liu, Q. Ma, J. Yuan, Y. Wang, M. Du, M. Liu, Effects of temperature and humidity on the daily new cases and new deaths of COVID-19 in 166 countries, Sci. Total Environ. 729 (2020). https://doi.org/10.1016/j.scitotenv.2020.139051.

[26] Q. Bukhari, Y. Jameel, Will coronavirus pandemic diminish by summer?, n.d. https://ssrn.com/abstract=3556998.

[27] J. Xu, Y. Zhang, How significant is a protein structure similarity with TM-score = 0.5?, Bioinformatics. 26 (2010) 889–895. https://doi.org/10.1093/bioinformatics/btq066.

[28] H.M. Berman, J. Westbrook, Z. Feng, G. Gilliland, T.N. Bhat, H. Weissig, I.N. Shindyalov, P.E. Bourne. The Protein Data Bank Nuc. Acids Res. 28 (2000) 235–242.

[29] A.C. Walls, M.A. Tortorici, B.J. Bosch, B. Frenz, P.J.M. Rottier, F. DiMaio, F.A. Rey, D. Veesler, Cryo-electron microscopy structure of a coronavirus spike glycoprotein trimer, Nature. 531 (2016) 114–117. https://doi.org/10.1038/nature16988.

[30] J. Guillén, A. J. Pérez-Berná, M. R. Moreno, J. Villalaín. Identification of the Membrane-Active Regions of the Severe Acute Respiratory Syndrome Coronavirus Spike Membrane Glycoprotein Using a 16/18-Mer Peptide Scan: Implications for the Viral Fusion Mechanism J. Virol. 79 (2005) 1743–1752. doi: 10.1128/JVI.79.3.1743-1752.2005

[31] J. Shang, Y. Wan, C. Liu, B. Yount, K. Gully, Y. Yang, A. Auerbach, G. Peng, R. Baric, F. Li, Structure of mouse coronavirus spike protein complexed with receptor reveals mechanism for viral entry, PLoS Pathog. 16 (2020). https://doi.org/10.1371/journal.ppat.1008392.

[32] C.M. Coleman, Y. V Liu, H. Mu, J.K. Taylor, M. Massare, D.C. Flyer, G.M. Glenn, G.E. Smith, M.B. Frieman, Purified coronavirus spike protein nanoparticles induce coronavirus neutralizing antibodies in mice, Vaccine 32 (2014) 3169–3174.

[33] L. Jiaming, Y. Yanfeng, D. Yao, H. Yawei, B. Linlin, H. Baoying, Y. Jinghua, G.F. Gao, Q. Chuan, T. Wenjie, The recombinant N-terminal domain of spike proteins is a potential vaccine against Middle East respiratory syndrome coronavirus (MERS-CoV) infection, Vaccine. 35 (2017) 10–18. https://doi.org/10.1016/j.vaccine.2016.11.064.

[34] J. Li, L. Ulitzky, E. Silberstein, D.R. Taylor, R. Viscidi, Immunogenicity and protection efficacy of monomeric and trimeric recombinant SARS coronavirus spike protein subunit vaccine candidates, Viral Immunol. 26 (2013) 126–132. https://doi.org/10.1089/vim.2012.0076.

[35] S. Zaim, J.H. Chong, V. Sankaranarayanan, A. Harky, COVID-19 and Multiorgan Response, Curr. Probl. Cardiol. (2020). https://doi.org/10.1016/j.cpcardiol.2020.100618.

[36] Y. He, Y. Zhou, S. Liu, Z. Kou, W. Li, M. Farzan, S. Jiang, Receptor-binding domain of SARS-CoV spike protein induces highly potent neutralizing antibodies: Implication for developing subunit vaccine, Biochem. Biophys. Res. Commun. 324 (2004) 773–781. https://doi.org/10.1016/j.bbrc.2004.09.106

[37] W. Tai, L. He, X. Zhang, J. Pu, D. Voronin, S. Jiang, Y. Zhou, L. Du, Characterization of the receptor-binding domain (RBD) of 2019 novel coronavirus: implication for development of RBD protein as a viral attachment inhibitor and vaccine, Cell. Mol. Immunol. (2020). https://doi.org/10.1038/s41423-020-0400-4.

[38] F. Li, Structure, Function, and Evolution of Coronavirus Spike Proteins, Annu. Rev. Virol. 3 (2016) 237–261. https://doi.org/10.1146/annurev-virology-110615-042301.

[39] F. Li, Receptor Recognition Mechanisms of Coronaviruses: a Decade of Structural Studies, J. Virol. 89 (2015) 1954–1964. https://doi.org/10.1128/jvi.02615-14.

[40] GROMACS development team, Gromacs documentation Release 2020. 1, (2020). http://manual.gromacs.org/documentation/5.1.4/%0Ahttp://files/1243/5.1.html.

[41] K. Lindorff-Larsen, S. Piana, K. Palmo, P. Maragakis, J.L. Klepeis, R.O. Dror, D.E. Shaw, Improved side-chain torsion potentials for the Amber ff99SB protein force field, Proteins Struct. Funct. Bioinforma. 78 (2010) 1950–1958. https://doi.org/10.1002/prot.22711.

[42] E.W. Sayers, T. Barrett, D.A. Benson, S.H. Bryant, K. Canese, V. Chetvernin., D.M. Church, M. Dicuccio, R. Edgar, S. Federhen, M. Feolo, L.Y. Geer, W. Helmberg, Y. Kapustin, D. Landsman, D.J. Lipman, T.L. Madden, D.R. Maglott, V. Miller, I. Mizrachi, J. Ostell, K.D. Pruitt, G.D. Schuler, E. Sequeira, S.T. Sherry, M. Shumway, K. Sirotkin, A. Souvorov, G. Starchenko, T.A. Tatusova, L. Wagner, E. Yaschenko, J. Ye, Database resources of the National Center for Biotechnology Information, Nucleic Acids Res. 37 (2009) 5–15. https://doi.org/10.1093/nar/gkn741.

[43] W. Humphrey, A. Dalke, and K. Schulten, VMD - Visual Molecular Dynamics J. Molec. Graphics 14 (1996) 33–38.

