## Supplementary material for "Investigation of the effect of temperature on the structure of SARS-Cov-2 Spike Protein by Molecular Dynamics Simulations": Supoorting Information

**Table S1. List of systems studied**

| Serial No. | Temperature (in ℃) | Simulation Time (ns) |
|---|---|---|
| 1 | 10 | 200 |
| 2 | 20 | 200 |
| 3 | 30 | 200 |
| 4 | 40 | 200 |
| 5 | 50 | 200 |
| 6 | 70 | 100 |

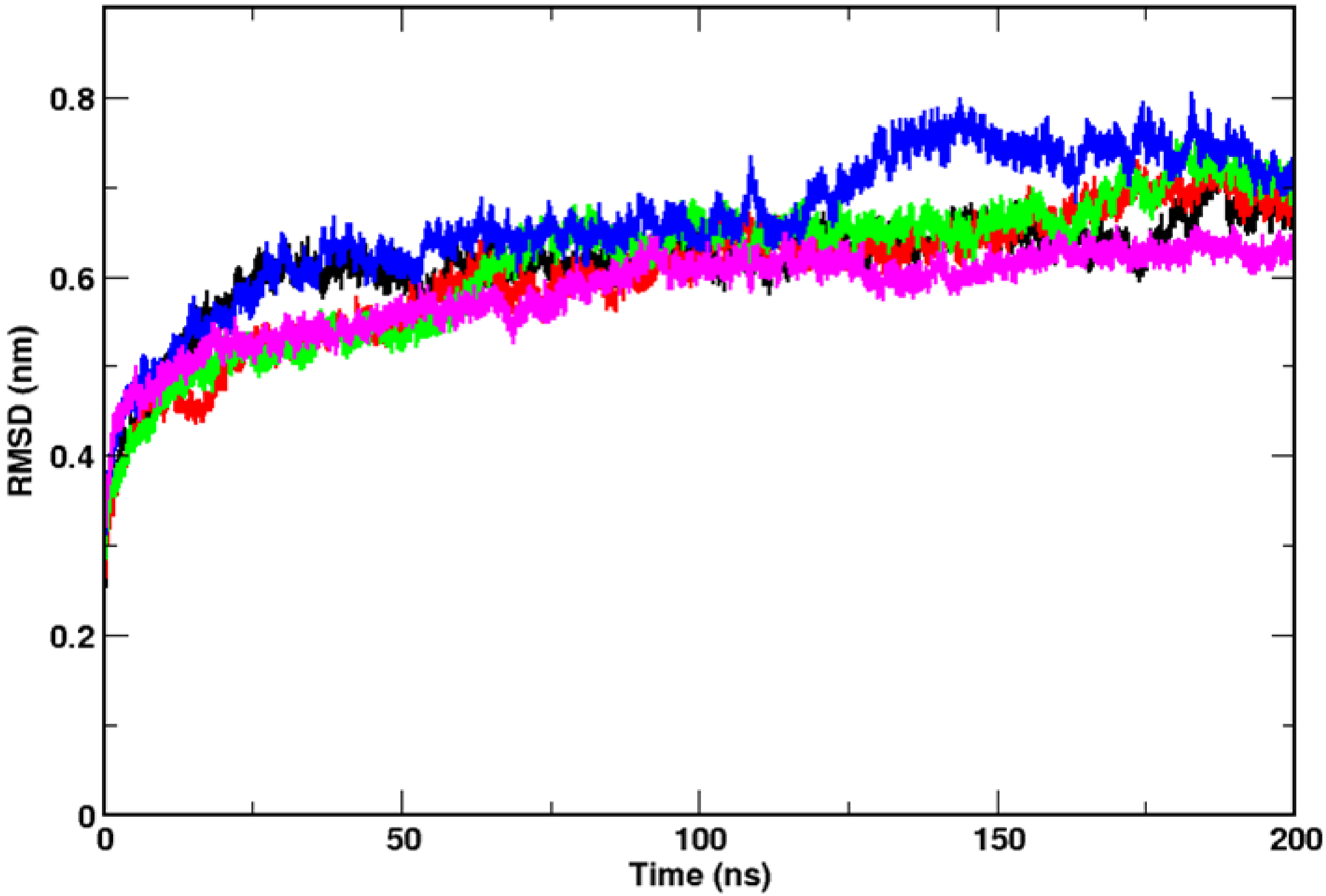


**Figure S1. Stability of the Spike protein across all temperatures.** RMSDs of Spike glycoprotein at 10 ℃(black), 20 ℃ (red), 30 ℃ (green), 40 ℃ (blue) and 50 ℃ (magenta) during 200ns of classical MD simulations showing stability of the simulations

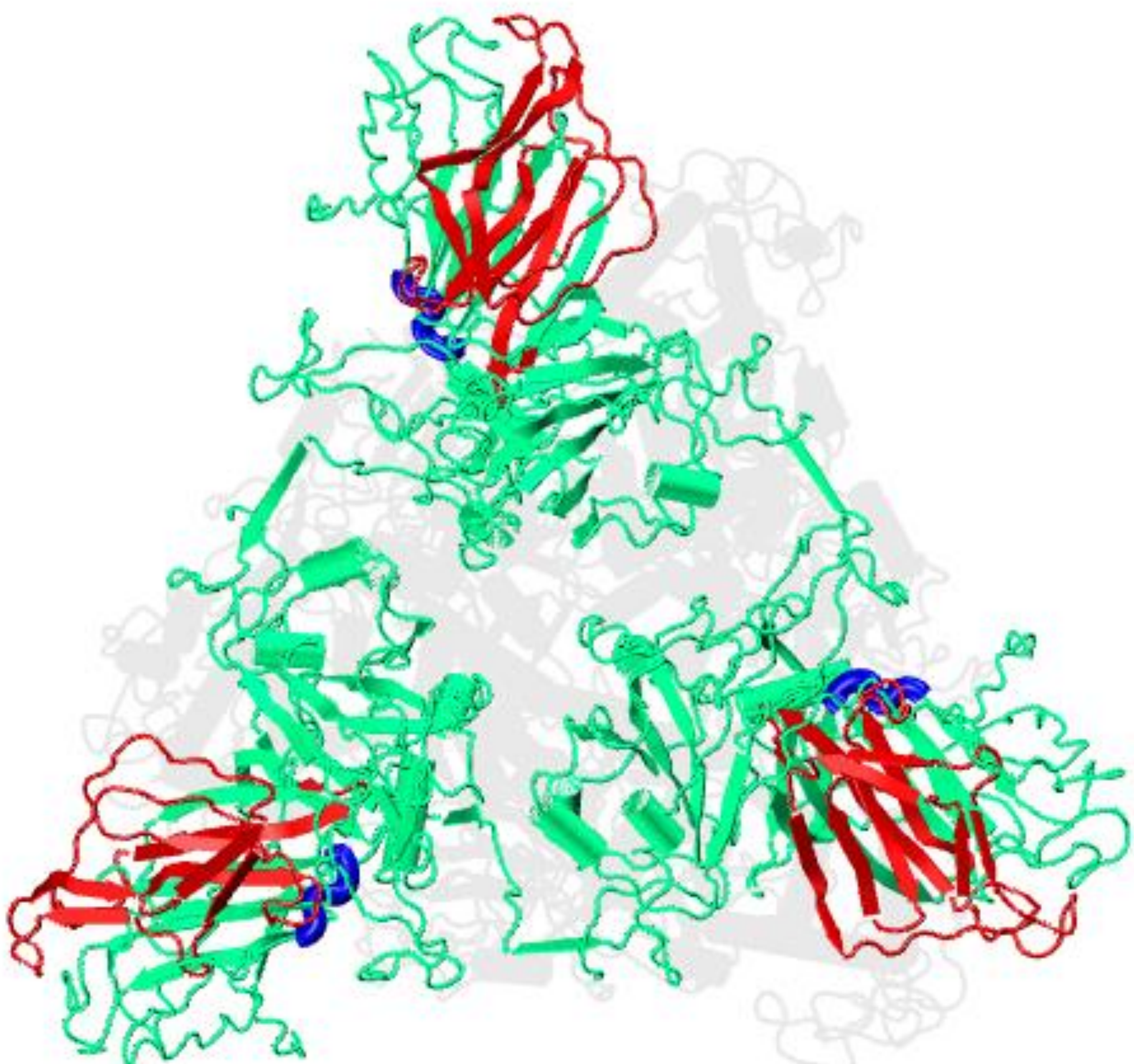

**Figure S2. Highly fluctuating regions in the N-Terminal Domain** Structure showing the S1 domain from top, highlighting the regions where peaks were observed in RMSF. The β4-β5 loop is shown in blue and the solvent exposed β6-β12 loop is shown in red

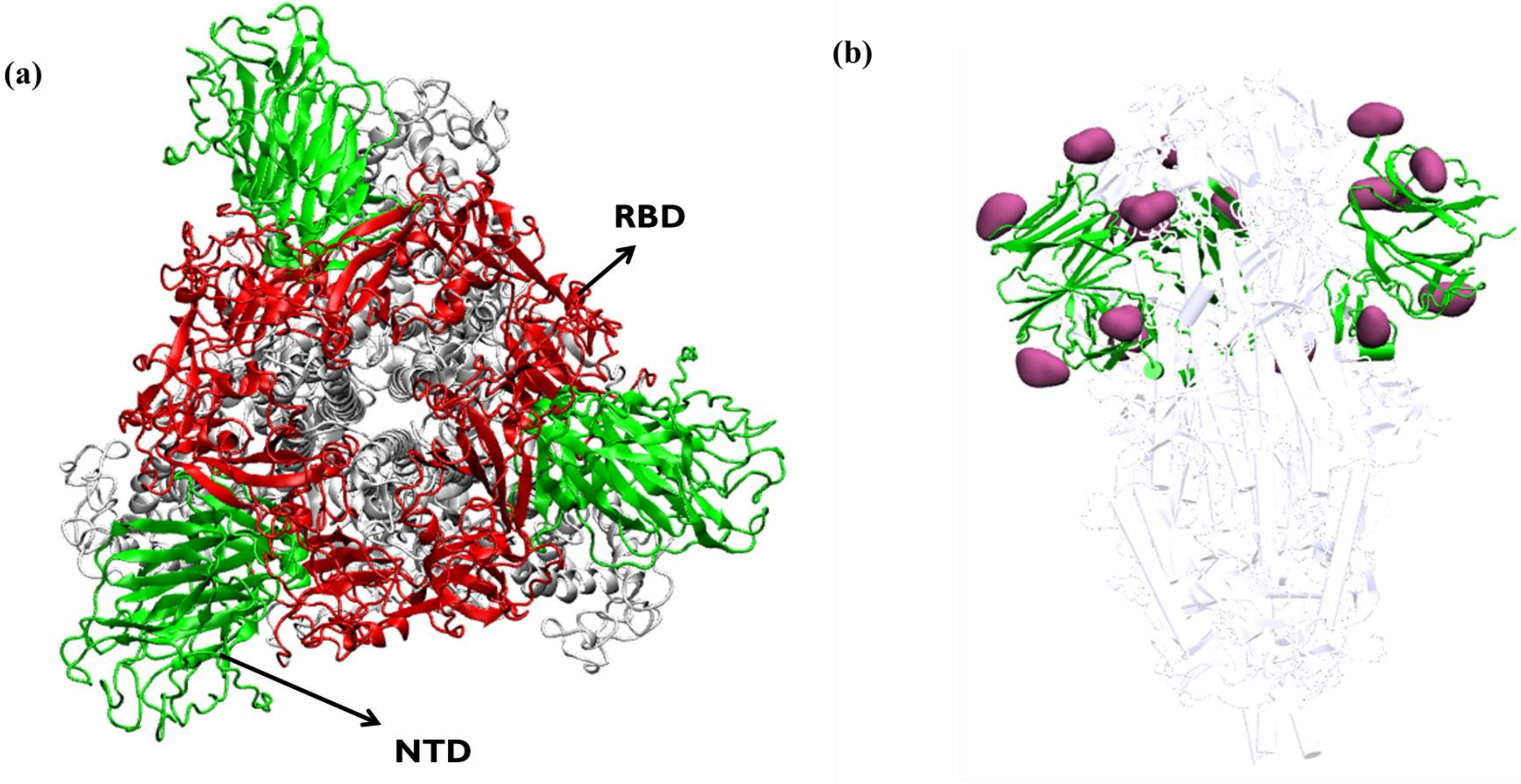


**Figure S3. The structure of S1 domain.** (a) Structure of S1 domain highlighting the N-Terminal Domain (in green) and Receptor Binding Domain (in red). (b) The orientation of N-acetyl glucosamine (NAG) residues around N-Terminal Domain in the crystal structure of Spike protein (PDB: 6VXX). The NAG residues are shown in surf mode and colored in pink, the N-Terminal Domain is shown in green

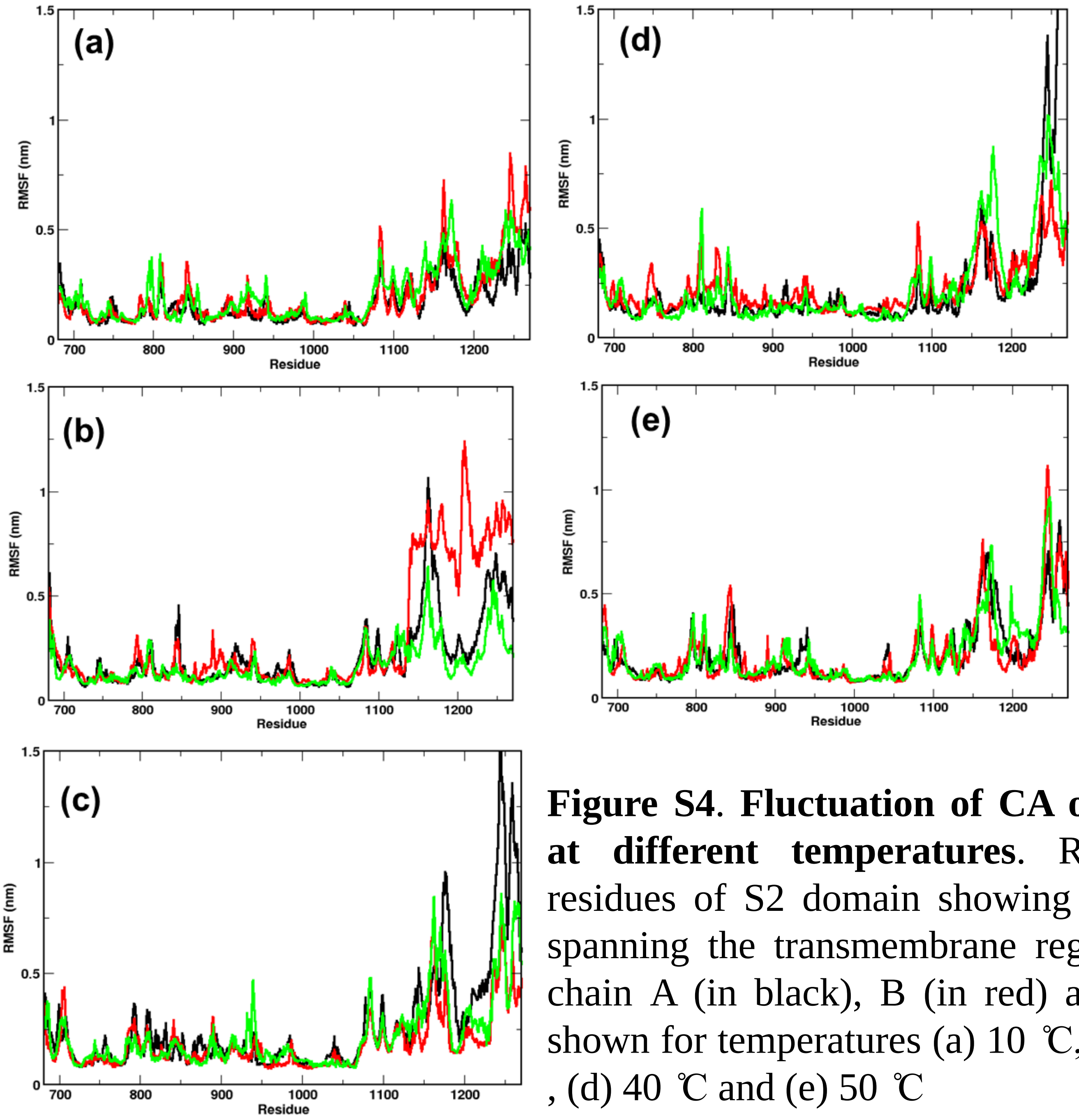


**Figure S4**. **Fluctuation of CA of individual chains at different temperatures**. RMSFs of the CA residues of S2 domain showing stability in residues spanning the transmembrane region. Fluctuations of chain A (in black), B (in red) and C (in green) are shown for temperatures (a) 10 ℃, (b) 20 ℃ , (c) 30 ℃ , (d) 40 ℃ and (e) 50 ℃

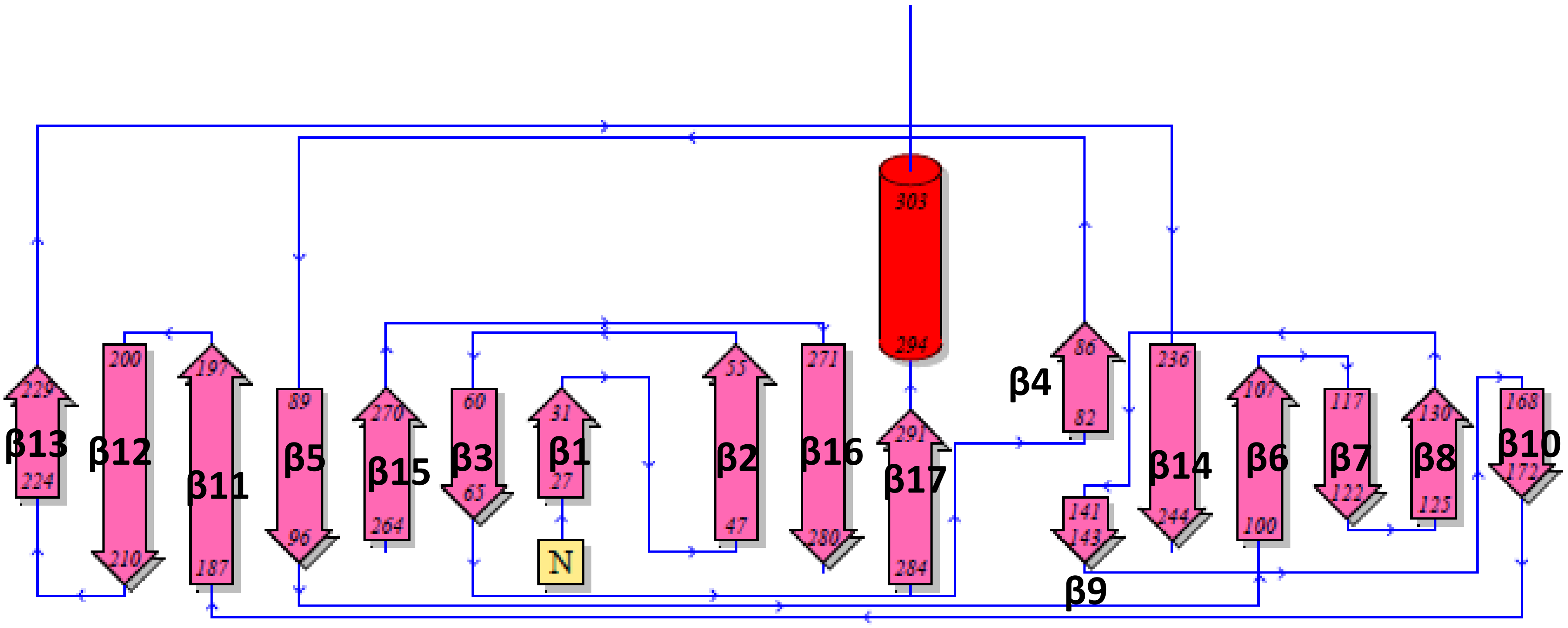


**Figure S5. Secondary structure of the N-Terminal Domain.** Topology of the N-Terminal Domain showing the three distinct layers of β sheet. β strands are colored in pink and α helix in red. The beginning and end residues are shown. The structure is generated from the pdbsum online server using the crystal structure of Spike protein (PDB ID: 6VXX)

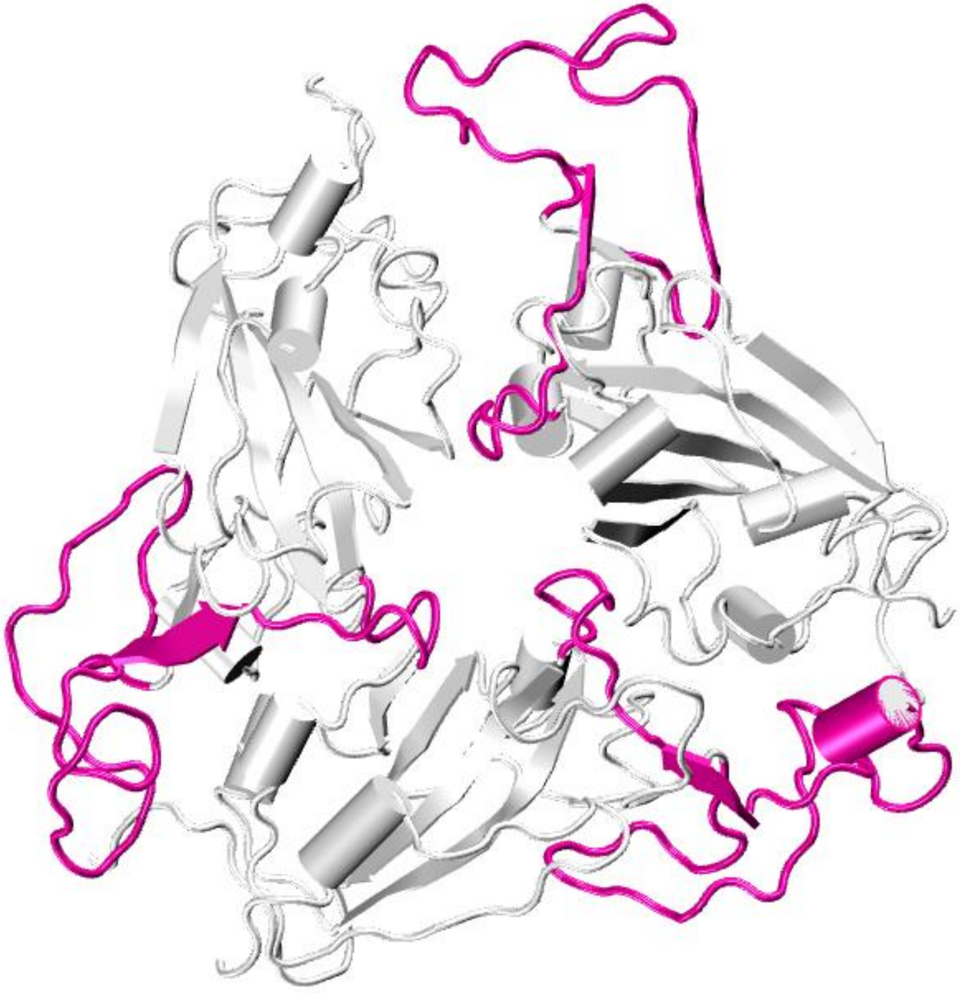

**Figure S6. Spike protein receptor binding motif at high temperature.** The time average conformation of the receptor binding motif highlighted in magenta at 70 ℃ after 100ns of simulation, showing the confined arrangement of loops
